# DRPBind: prediction of DNA, RNA and protein binding residues in intrinsically disordered protein sequences

**DOI:** 10.1101/2023.03.20.533427

**Authors:** Ronesh Sharma, Tatsuhiko Tsunoda, Alok Sharma

## Abstract

DRPbind predicts three components: deoxyribonucleic acid (DNA) binding, ribonucleic acid (RNA) binding and protein-binding residues of query protein sequences. DRPbind utilizes independent sources of information encoded for each component predictor and relies on the information-rich profiles of protein evolutionary-based features, protein physicochemical-based features, and protein structural-based features. DRPbind employs protein profile-based features extracted from the bigrams of PSSM and HMM profiles. It also extracts features from physicochemical and structural attributes. DRPbind takes primary protein sequences as input, and through the Support Vector Machine (SVM) classifier, it provides the binding prediction. DRPbind is optimized based on a specific binding type and shown superior performance in terms of simultaneously predicting the DNA-binding, RNA-binding and protein-binding residues. The source code is available at https://github.com/roneshsharma/DNA-RNA-Protein-Binding/wiki

## Background

Proteins with intrinsically disordered regions (IDRs) play a crucial role in various biological processes [1–3]. Many IDRs interact with nucleic acids and proteins [4–6]. Annotation of these IDRs provides biological insight and understanding of protein function. Computational prediction of IDRs in the protein sequence is a challenging task. In this study, we present the DRPbind predictor to predict the deoxyribonucleic acid (DNA), ribonucleic acid (RNA) and protein-binding residues in the protein sequence. These binding residues in the protein sequence interact with many different complexes and are associated with important diseases. Identifying the protein binding residues accurately is a step towards understanding the gene expression and supporting the development of precision medicine. Wet lab experimental methods allow protein binding residues to be analysed at the molecular level; however, these experimental approaches are expensive and time-consuming. Accordingly, there is a demand to introduce efficient computational methods to identify the protein binding residues.

Despite the recent efforts to improve the performance of the DNA, RNA and protein-binding predictors, the prediction accuracy remains limited. In this study, we propose DRPbind to predict the DNA, RNA and protein-binding residues. To build this predictor, we first extract PSSM and HMM profiles of the protein sequence using PSI-blast [7] and HHblits [8] tools, respectively. Then we obtain structural attributes of the protein sequence using the spider2 tool [9] and we employ physicochemical indexes to represent the protein sequence. Next, we extract features, including amino acid composition (AAC), bigram [10], and auto-covariance (AC), from the protein sequence. To build DRPbind, the extracted feature vector is fed to the Support Vector Machine (SVM) classifier to predict the binding residues. We have also conceptualized an extension of DRPbind, DeepDRPbind. The predictor DeepDRPbind is not analyzed in this paper, however, this predictor is employed by the CAID-2 project, where many published and unpublished predictors are compared (https://predictioncenter.org/casp15/). The concept of DeepDRPbind is developed by tensor flow of the features from the protein sequences. The extracted features are converted to corresponding well-organized images in a unique way through the application of the DeepInsight tool [11]. The obtained features in tabular form are transformed into images which enables the application of convolutional neural networks (CNNs) for the binding residue prediction. In this paper, our results demonstrate that DRPbind enhances binding prediction and gives promising results compared to the state-of-the-art method.

## Method

We employ the data sets previously introduced by Zhang et al. [4]. Table 1 shows the details of the data set containing training, validation and test sets.

**Table 1:** Benchmark dataset

| Dataset | Number of proteins | Number of disordered residues |  |  |  |
| --- | --- | --- | --- | --- | --- |
|  |  | Protein-binding | DNA-binding | RNA-binding | All disordered |
| Training | 238 | 15,341 | 2913 | 1437 | 27304 |
| Validation | 118 | 6,464 | 1284 | 608 | 11716 |
| Test | 394 | 17,540 | 2377 | 1518 | 46041 |

In this study, we develop DRPbind predictor to predict the DNA, RNA and protein-binding residues (Figure 1). First, we extract PSSM and HMM profiles using PSI-blast [7] and HHblits [8] tools. Then, we extract the structural attributes using the spider2 [9] tool and employ physicochemical attributes to represent the amino acids of the protein sequences [12]. Next, we compute the features, including AAC, bigram [10] and AC [13] from the extracted profiles and attributes. The reason is to integrate various attributes representing a protein sequence for binding predictions. After that, an SVM classifier is tuned and applied to the extracted features to build the DRPbind predictor. Figure 1 also depicts the DeepDRPbind predictor. However, since its results are not discussed in this paper, it has not been described in detail. Nonetheless, the DeepDRPbind predictor has shown superior performance for protein binding site prediction when compared by independent assessors using 33 methods (see pages 27 and 28 of CAID-2 project report, CASP15 conference, 10-13 December 2022, Antalya): *https://predictioncenter.org/casp15/doc/presentations/Day1/DPiovesan_CAID2.pdf*.

**Figure 1:**
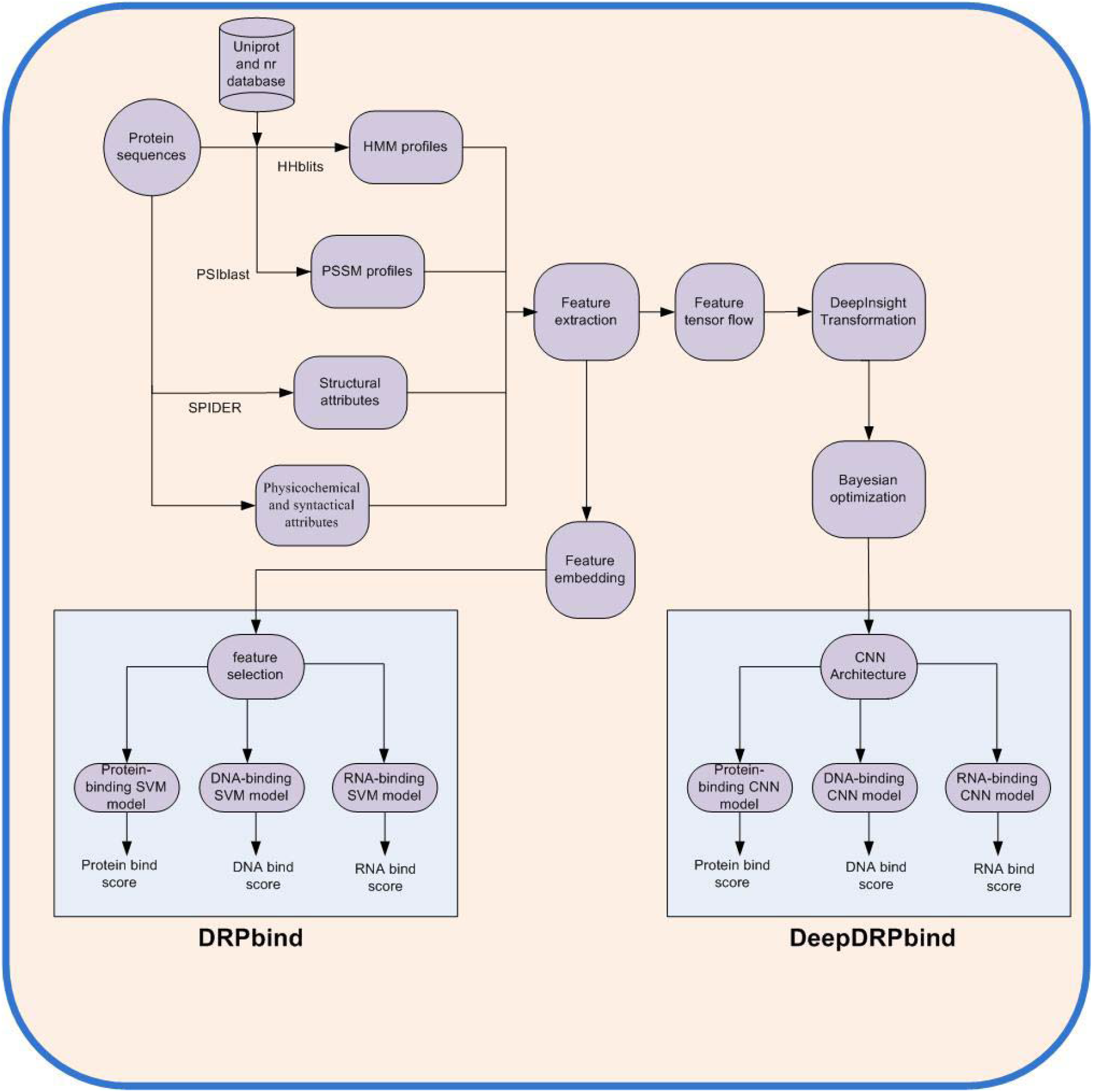
Proposed overview of the system.

### A. Feature extraction method

A comprehensive set of features are extracted to characterize a protein sequence. In terms of evolutionary information, PSSM and HMM profiles are generated using PSI-blast [7] and HHblits [8] tools, respectively. PSI-blast produces PSSM profiles of size ***L*** by 20 and HHblits produces HMM profiles of size ***L*** by 30, where ***L*** is the length of the protein sequence. To produce these profiles, PSI-blast and HHblits iteratively search through the protein databases (nr and uniprot20) and build multiple sequence alignments (MSAs) from which protein profiles are computed to represent the 20 standard amino acids in the homologous protein. Compared to the PSSM profiles, the HMM profiles contain additional information describing the insertion, deletion and match during MSAs. However, for this study, we only use the first 20 columns. For physicochemical properties, we have used the physicochemical properties included in standard 544 amino acid indexes [12], these indexes are available at ftp://ftp.genome.jp/pub/db/community/aaindex/. The set contains 27 amino acid indexes. These indexes have shown promising results for molecular recognition feature (MoRF) [14] prediction.

Furthermore, for structural information, the spider2 tool [9] is used to generate the structural attributes, including secondary structure states (SSS), solvent accessible surface area (ASA), backbone angles and half-sphere exposure (HSE). SSS represents the description of each amino acid in several discrete states, such as helix, sheet and coil, ASA measures the exposure level of amino acid to solvent in a protein sequence, backbone angles contain backbone dihedral angles of amino acids in the protein sequence and HSE is an alternative measure of the solvent exposure of amino acid.

Using the above profiles, properties and attributes, features including AAC, bigram and AC, are computed as follows:

1. AAC: this feature is computed from the profiles of the protein sequences. Let matrix ***H*** of size ***L×C*** be the profile of a protein sequence, where ***L*** is the length of the protein sequence and ***C*** refers to the number of attributes/columns. The computation of the feature vector from the matrix *H* is as follows:

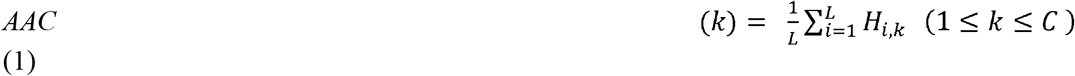

where *H_i,k_* is the element of the profile matrix. Computing *AAC*(*k*) for *k* ranging from 1 to *C* would give a feature vector of dimension *C*.
2. Bigram: this feature is computed from the matrix *H* as follows:

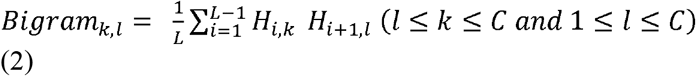 Computing *Bigram_k,l_* for *k* = 1,2, …., *C* and *I* = 1,2,….,*C* would give a matrix of size *C* × *C*. This matrix can be represented in a vector form by reshaping the *C* × *C* matrix into a vector of length *C*^2^.
3. AC: this feature is computed from the matrix *H* as follows:

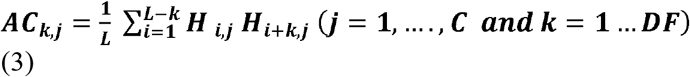

where ***DF*** is the distance factor and computing the ***AC*** frequencies for ***j*** = **1**,….,***C*** and ***k*** = **1**… ***DF*** would give an ***AC*** matrix of size ***DF × C***. This matrix can be represented in a vector form by reshaping ***DF × C*** matrix into a vector of fixed length.

### B. Experimentation

To build and evaluate the proposed method, we employed SVM and CNN classifiers for binding residue prediction. To conduct the experiment, MATLAB software is utilized with the LibSVM [15] package adopted with radial basis function (RBF) kernel, and a grid search is performed to select the kernel parameters. Moreover, Bayesian optimization is adopted to tune the CNN network to predict the binding residues. The combined training and validation set is used to train the model, while the test set is used to evaluate and validate the performance of the proposed method.

## Results and Discussion

DRPbind is evaluated on a benchmark dataset that has been previously utilised to evaluate DNA, RNA and protein binding predictors. The result (Table 2) demonstrates that DRPbind outperforms the state-of-the-art predictor (DeepDISOBind [4]) and is the most accurate predictor in terms of simultaneously predicting the DNA, RNA and protein-binding residues.

**Table 2:** Result comparison

| Predictors | Protein-binding | DNA-binding | RNA-binding |
| --- | --- | --- | --- |
| DeepDISOBind | 0.771 | 0.746 | 0.736 |
| DRPBind (proposed) | 0.765 | 0.780 | 0.747 |

## Conclusions

In conclusion, DRPbind has been shown to be effective in predicting the binding residues of DNA, RNA, and proteins in query protein sequences. This method utilized various sources of information, including evolutionary-based features, physicochemical-based features, and structural-based features, and employed SVM for prediction. Overall, the optimized performance of DRPbind in predicting multiple types of binding residues highlights their potential as valuable tools for protein structure and function analysis.

## Declarations

### Competing interests

We have no competing interests.

### Authors’ contributions

RS and AS conceived the project and processed the data. RS and AS performed the analysis and wrote the manuscript. TT supervised and assisted in the manuscript writeup. All authors read and approved the final manuscript.

## Notes

https://github.com/roneshsharma/DNA-RNA-Protein-Binding/wiki

### Competing Interest Statement

The authors have declared no competing interest.

